# Cell Stiffening Contributes to Complement-mediated Injury of Choroidal Endothelial Cells in Early AMD

**DOI:** 10.1101/2021.10.06.463274

**Authors:** Andrea P. Cabrera, Jonathan Stoddard, Irene Santiago Tierno, Nikolaos Matisioudis, Mahesh Agarwal, Lauren Renner, Neha Palegar, Martha Neuringer, Trevor McGill, Kaustabh Ghosh

## Abstract

Age-related macular degeneration (AMD) is the leading cause of blindness in the aging population. Yet, no therapies exist for ∼85% of all AMD patients who have the dry form that is marked by degeneration of the retinal pigment epithelium (RPE) and underlying choroidal vasculature. As the choroidal vessels are crucial for RPE development and maintenance, understanding how they degenerate may lead to effective therapies for dry AMD. One likely causative factor for choroidal vascular loss is the cytolytic membrane attack complex (MAC) of the complement pathway that is abundant on choroidal vessels of humans with early dry AMD. To examine this possibility, we studied the effect of complement activation on choroidal endothelial cells (ECs) isolated from a rhesus monkey model of early AMD that, we report, exhibits MAC deposition and choriocapillaris endothelial loss similar to that seen in human early AMD. Treatment of choroidal ECs from AMD eyes with complement-competent normal human serum caused extensive actin cytoskeletal injury that was significantly less pronounced in choroidal ECs from young normal monkey eyes. We further show that ECs from AMD eyes are significantly stiffer than their younger counterparts and exhibit peripheral actin organization that is distinct from the longitudinal stress fibers in young ECs. Finally, these differences in complement susceptibility and mechanostructural properties were found to be regulated by the differential activity of small GTPases Rac and Rho because Rac inhibition in AMD cells led to simultaneous reduction in stiffness and complement susceptibility while Rho inhibition in young cells exacerbated complement injury. Thus, by identifying cell stiffness and cytoskeletal regulators Rac and Rho as important determinants of complement susceptibility, the current findings offer a new mechanistic insight into choroidal vascular loss in early AMD that warrants further investigation for assessment of translational potential.

## Introduction

Age-related macular degeneration (AMD) is the leading cause of vision loss in the elderly population. Yet, no therapies exist for almost 85% of all AMD patients who have the dry form of this disease [1,2]. Since early dry AMD is a potential risk factor for the vision-threatening advanced forms of geographic atrophy (‘dry’) or neovascular (‘wet’) AMD, there is significant interest in understanding and inhibiting AMD pathogenesis in the early dry stage [2]. Early AMD is characterized by dysfunction and atrophy of retinal pigment epithelium (RPE) and the underlying choriocapillaris that, together, are implicated in the accumulation of lipid-rich drusen deposits in the sub-RPE space [3,4]. In addition to RPE abnormalities, the loss of choriocapillaris (in early AMD) and choroidal vessels (in advanced geographic atrophy) is being increasingly recognized as a potentially crucial factor in AMD pathogenesis [5]. This is because firstly, choroidal vessels provide vital metabolic support to the overlying RPE and, thus, their loss is expected to cause hypoxia-related RPE dysfunction, and secondly, new studies reveal that choroidal endothelial cells (ECs) provide instructive cues for RPE maturation and function, as well as choroidal immunomodulation [6,7]. This dependence of RPE on choroidal vessels provides the rationale to investigate how the latter degenerates in early AMD.

The complement system, an integral component of the innate immune response, is dysregulated in AMD, leading to the deposition of membrane attack complex (MAC; C5b-9) on choroidal vessels of humans with early AMD [8,9]. Since cultured ECs subjected to complement activation undergo membrane MAC deposition and injury [10,11], it is possible that MAC deposition on choroidal vessels contributes to their degeneration in early AMD. However, MAC is also found on the healthy choroidal vessels of young normal human eyes [8]. This unexpected observation raises the possibility that specific age-related factors promote MAC-induced choroidal vascular atrophy in early AMD.

Findings from our recent studies using a replicative senescence model of aging support this view. Specifically, we found that senescence leads to an increase in chorioretinal RF/6A cell stiffness that, in turn, exacerbates MAC injury by altering cellular mechanotransduction, the process by which mechanical cues are converted into biochemical signaling pathways that, ultimately, govern cell behavior [10,12,13]. Although this study is the first to implicate an age-related factor, viz. senescence-associated cell stiffening and mechanotransduction, in complement injury, it does not accurately reflect the pathogenesis of choroidal vascular loss in early AMD. This is because firstly, senescence is not unique to aging and is one of many changes that occur in biologically aging cells and tissues [14], and secondly, immortalized cell lines do not fully recapitulate the molecular and functional characteristics of primary cells that constitute aging or diseased tissues.

Thus, to enhance the translational impact of our mechanistic understanding of choroidal vascular loss, we here studied the effect of complement activation on choroidal ECs isolated from a clinically relevant rhesus monkey (*Macaca mulatta*) model of early AMD. Rhesus monkeys are a unique model for the study of early AMD pathogenesis because, unlike non-primate models, they (a) have a retinal structure closely resembling that found in humans, (b) share the same AMD susceptibility genes *ARMS2* and *HTRA1* with humans, and (c) undergo drusen accumulation as seen in humans [15]. Using this model, we show that, when compared with choroidal ECs isolated from young normal monkey eyes, those obtained from old drusen-laden eyes exhibit a distinct peripheral actin organization, are significantly stiffer, and undergo greater MAC injury, attributes that can be significantly reversed by modulating the levels of cytoskeletal mediators of mechanotransduction Rho and Rac.

## Materials and Methods

### Retinal Fundus Imaging

Retinal fundus color images were obtained from rhesus monkeys using a standard fundus camera (Zeiss FF450). Photographs were graded and categorized based on the presence, number, and size of drusen, as previous reported [16]. For these studies, three groups were selected: young normal with no drusen (YN, 6-9 years old), old normal with no drusen (ON, 14-21 years old), and old with moderate/severe drusen (OD, 14-30 years old).

### Immunolabeling and Image Analysis of Retinal Sections

Rhesus monkey eyes were obtained through the ONPRC Tissue Distribution Program within 10 minutes of humane euthanasia. Eyes selected for immunolabeling (YN, 7-9 years old, 2 female, 1 male; ON, 14-21 years old, 2 female, 1 male; and OD, 14-30 years old, 2 female, 1 male) were immersion-fixed in 4% paraformaldehyde for 24h, cryoprotected using 10%, 20%, and 30% sucrose concentrations and frozen in Tissue-Tek O.C.T. Compound (Sakura, Japan). Cryosections cut at 14μm-thick were labeled with C5b-9 and collagen IV antibodies (AbCam), counterstained with DAPI followed by autofluorescence eliminator (Millipore). Mounted sections were imaged using a Zeiss LSM 880 (with Airyscan) confocal microscope.

To quantify MAC (C5b-9) intensity, ImageJ was used to select the choriocapillaris layer (from the collagen IV-labeled image) as the region of interest (ROI) before applying an ROI binary mask to the anti-C5b-9-labeled image. Mean MAC (C5b-9) fluorescence intensity was then measured from at least 90 images from six retinal sections and normalized per unit area. Quantification of acellular capillaries was performed by first superimposing the collagen IV-labeled image of choriocapillaris on its corresponding DAPI-labeled image, followed by counting the number of capillaries that lacked DAPI-labeled cell nuclei on the luminal side of collagen IV-positive capillaries and dividing it by the total number of capillaries. At least 70 images from five retinal sections were analyzed.

### Cell Isolation

Macular choroidal endothelial cells (ECs) were isolated from three groups of female rhesus monkeys (YN, 6-9 years old; ON, 14-20 years old; OD, 18-19 years old) using a previously reported protocol [17] that is described in detail in *Supplementary Material*. Also described in *Supplementary Material* is the isolation of monkey retinal pigment epithelial (RPE) cells for use as a positive control in the detection of contaminating (RPE65-expressing) RPE cells in choroidal EC cultures.

### Cell Culture

Monkey choroidal ECs between passages 4-9 were grown on 0.5% (w/v) gelatin-coated dishes in MCDB 131 basal medium supplemented with 10% FBS, 10 mM L-glutamine (Life Technologies), 10 ng/mL EGF, 4 ng/mL bFGF, 1 ug/mL hydrocortisone (Sigma), and 1x antibiotic/antimycotic. EC cultures seeded at 10,000 cells/cm^2^ were maintained at 37°C with 5% CO_2_/95% humidity and passaged every two days (80-90% confluence). For all in vitro studies, cells were seeded at 40,000 cells/cm^2^ in starvation medium (MCDB 131 basal medium supplemented with 2.5% FBS and 1x antibiotic/antimycotic) for 6h prior to assays.

### Quantitative RT-PCR

Total RNA was obtained from choroidal EC cultures and frozen RPE cells using an RNA purification kit (Direct-zol RNA MiniPrep; Zymo Research, Irvine, CA, USA), as per the manufacturer’s protocol. Isolated RNA was converted to cDNA with high capacity cDNA reverse transcription kit (Applied Biosystems), and amplified with TaqMan PCR primers for VE-cadherin, VEGFR2, or RPE65 (Thermo Fisher Scientific, Inc.) using the CFX connect real-time PCR detection system (BioRad, Hercules, CA, USA). Relative mRNA levels were determined by the comparative cycle threshold method with normalization to glyceraldehyde 3-phosphate dehydrogenase (GAPDH; Thermo Fisher Scientific, Inc.).

### Flow Cytometry

EC monolayer was detached with accutase (Corning) and the resulting single-cell suspension was labeled with Phycoerythrin (PE)-conjugated mouse anti-human CD146 antibody (BD Biosciences) or CD31 antibody (Abcam) for 20 min at 4°C in FACS buffer (0.5% BSA and 2mM EDTA). PE-conjugated mouse IgG was used as Isotype control. Approximately 25,000 cells were assessed on a NovoCyte flow cytometer (ACEA Biosciences, San Diego, CA) for CD146 or CD31 expression followed by quantitative analysis using FlowJo (Treestar, Inc., Ashland, OR, USA).

### Capillary Network Formation Assay

Cultrex^®^ basement membrane solution (10 mg/mL; Trevigen Inc.) was mixed with fibrin gel solution (2 mg/mL fibrinogen and 0.66 U/mL thrombin; Sigma) at a 1:1 (v/v) ratio and allowed to solidify for 4h at 37°C. Next, choroidal ECs were plated at 40,000 cells/cm^2^ on the surface of the three-dimensional hydrogel and cultured in growth medium supplemented with VEGF (25 ng/mL) and bFGF (30 ng/mL) for 6h prior to imaging with Nikon Eclipse TI epifluorescence microscope.

### Complement Activation and Injury Assay

ECs grown on gelatin-coated glutaraldehyde-crosslinked glass coverslips were treated with 5% complement-competent normal human serum (NHS; Complement Technology, Inc., Tyler, TX, USA) in veronal buffered saline (VBS; 145 mM NaCl, 1.8 mM sodium barbital, 3 mM barbituric acid, 50 mM CaCl_2_, and 25 mM MgCl_2_, pH 7.4) for 3h at 37°C to promote activation of the complement system. Next, untreated and NHS-treated cells were fixed with 4% PFA, incubated (without detergent-based permeabilization) with AlexaFluor 594 Phalloidin (Invitrogen; 20 min, RT) to visualize actin cytoskeleton, and imaged with Nikon Eclipse TI epifluorescence microscope. To assess cellular injury, ImageJ was used to quantify the percent cells with damaged actin stress fibers (n ≥ 60 cells per condition).

To determine the effects of Rho/Rac on complement injury, cells were co-treated with 5% NHS and either NSC23776 (NSC; 1 mM), a pharmacological Rac1 inhibitor, or Y27632 (Y27; 100 µM), a pharmacological inhibitor of Rho/Rho-associated kinase (ROCK), for 3h at 37°C.

### MAC (C5b-9) Immunolabeling of Cultured ECs

NHS-treated ECs were fixed in 4% PFA, blocked with 2% BSA, and incubated with anti-C5b-9 (AbCam) for 2h at RT followed by secondary antibody incubation for 1h at RT in the dark. Labeled cells were imaged with Zeiss LSM 880 (with Airyscan) confocal microscope. ImageJ was used to quantify net C5b-9 fluorescence intensity (n ≥ 60 cells per condition).

### Cell Stiffness

ECs were grown to confluence on glutaraldehyde-crosslinked gelatin-coated cover slips before replacing growth medium with calcium buffer (136 mM NaCl, 4.6 mM KCl, 1.2 mM MgSO_4_, 1.1 mM CaCl_2_, 1.2 mM KHPO_4_, 5mM NaHCO_3_, 5.5 mM glucose, and 20 mM HEPES with 2.5% FBS) prior to stiffness measurement using an MFP3D atomic force microscope (AFM; Asylum Research) in contact mode that applied a 2.5 nN indentation force on EC monolayers with a silicon nitride cantilever (0.1 N/m spring constant) fitted with a 5 μm diameter glass bead (Novascan). EC monolayers were indented at ≥20 distinct spots, with each spot indented three times to obtain average stiffness. For some studies, OD ECs were treated with Rac inhibitor NSC23776 (1 mM) for 30 min at 37°C prior to AFM indentation.

To examine the effect of complement injury on cell stiffness, untreated ECs (in calcium buffer containing 0.5% FBS) and NHS-treated ECs (2-3h in VBS buffer) were subjected to force indentation (0.2 nN set point) using a NanoWizard® 4 XP BioScience AFM (Bruker, CA, USA) fitted with a pre-calibrated PFQNM-LC-A-CAL probe (0.075 N/m spring constant) containing a 70 nm-radius hemispherical tip (Bruker AFM Probes, CA, USA). 15 cells were indented, with each cell indented at three distinct spots to obtain average stiffness.

### Rho and Rac Activity

Activities of RhoA and Rac1 GTPases were measured using the RhoA and Rac1 G-LISA activation assay kits, respectively (Cytoskeleton Inc, Denver, CO, USA), as per the manufacturer’s protocol.

### Western Blot

Confluent EC monolayers were lysed in RIPA lysis buffer (G-Biosciences, St. Louis, MO) containing protease and phosphatase inhibitors (Boston BioProducts, Ashland, MA), followed by centrifugation and loading of the equal amounts of protein into a 10% SDS-polyacrylamide gel. The separated proteins were transferred onto a nitrocellulose membrane for detection with polyclonal rabbit anti-Phospho-Myosin Light Chain 2 (Ser19) antibody (Cell Signaling Technologies) or monoclonal rabbit anti-Myosin Light Chain 2 (D18E2) antibody (Cell Signaling Technologies). GAPDH (Sigma) was used as the loading control. Protein bands were visualized using a chemiluminescent detection kit (Thermo Scientific) coupled with a camera-based imaging system (Biospectrum AC ImagingSystem). Densitometric analysis was performed by ImageJ.

### Statistics

All data were obtained from multiple cells and multiple replicates/condition and expressed as mean ± SEM or SD (as indicated in each respective section). Statistical significance was determined using analysis of variance (ANOVA), followed by Tukey’s and Bonferroni post-hoc analysis (Instat; GraphPad Software Inc., La Jolla, CA, USA). Results were considered significant if P< 0.05.

## Results

### Rhesus Monkey Eyes Exhibit Complement Activation and Choroidal Vascular Loss Associated with Early AMD

To determine the suitability of rhesus monkeys for the study of complement-mediated choroidal vascular atrophy seen in early AMD, we first classified the monkeys as young normal with no drusen (YN; 6-9 years old), old normal with no drusen (ON; 14-21 years) and old AMD monkeys with moderate-to-severe drusen (OD; 14-30 years) based on age and appearance of yellowish-white deposits in fundus photographs (Fig. 1A). Subsequent c5b-9 immunolabeling of retinal sections from each group revealed notable MAC deposition in the Bruch’s membrane and choriocapillaris that was significantly more pronounced in OD eyes (∼1.5-fold greater; p<0.001) than in YN eyes (Fig. 1B), a trend consistent with observations in humans [8]. Importantly, increased MAC deposition in the choriocapillaris of OD eyes was associated with significantly greater endothelial loss in these eyes (median value of 10%; p<0.001) than in YN or ON eyes (median value of 0%) (Fig. 1C).

**Figure 1:**
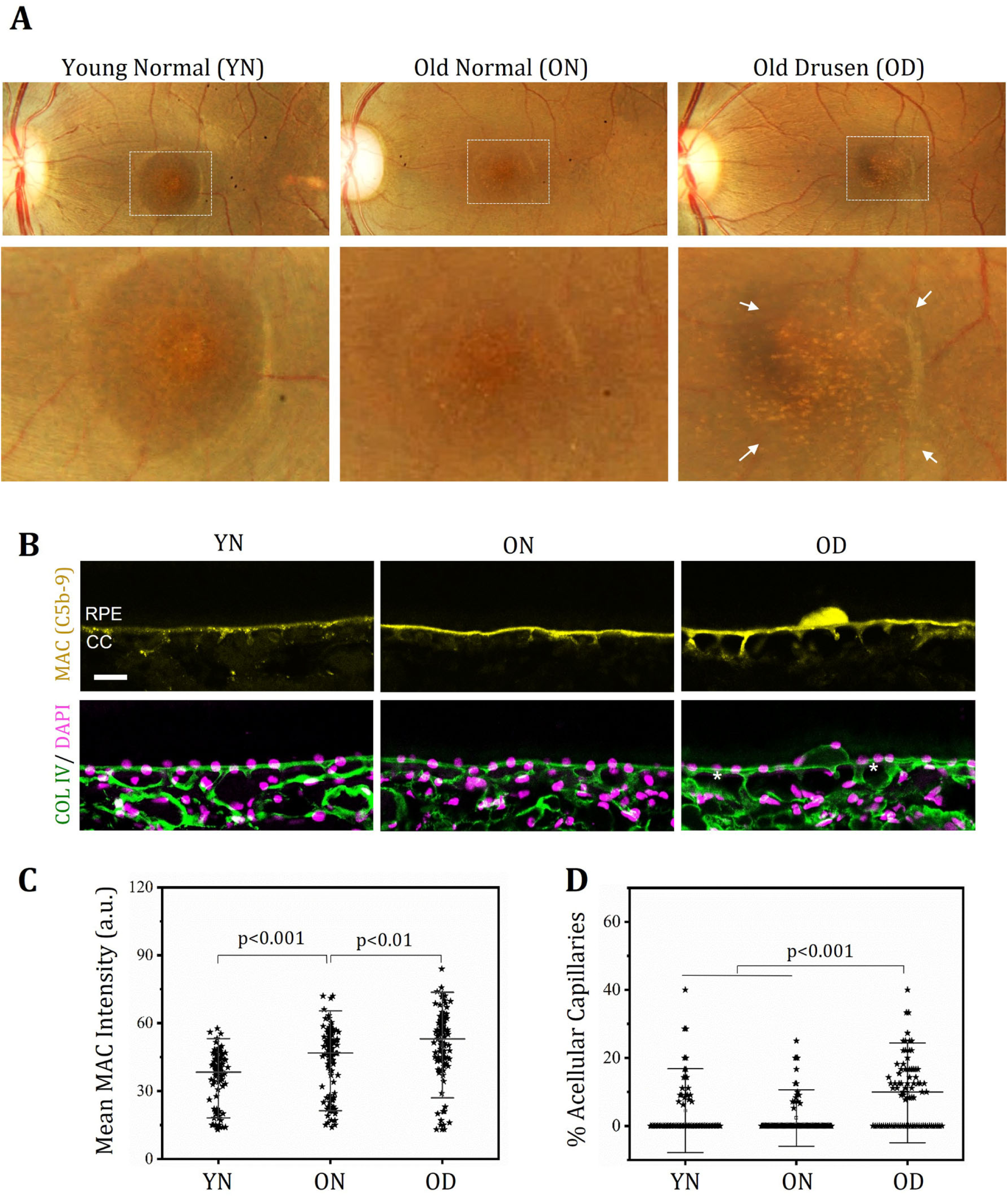
Rhesus monkeys with macular drusen exhibit increased MAC (C5b-9) deposition and choriocapillaris atrophy. **(A)** Retinal fundus images were used to categorize rhesus monkey eyes as young normal (YN; 6-9 years old), old normal (ON; 14-21 years old), or old with moderate-to-severe macular drusen (OD; 14-30 years old). *Bottom panel* shows magnified view of the macular region outlined in the *top panel*, with arrows indicating drusen accumulation in OD eyes. **(B)** Confocal images of macular sections from YN, ON, and OD eyes labeled with anti-C5b-9 (yellow) and anti-collagen IV (green) reveal MAC (C5b-9) immunoreactivity within drusen, and at the level of sub-RPE and choriocapillaris. Notably, MAC immunoreactivity was not observed in RPE. Acellular capillaries (asterisk) were identified by the absence of DAPI-labeled (purple) cell nucleus on the luminal side of collagen IV-positive capillaries. Scale bar 25 um. RPE, Retinal pigment epithelium; CC, choriocapillaris. **(C)** Fluorescence intensity analysis showed that OD eyes exhibit significantly higher (∼1.5-fold; p<0.001) MAC immunoreactivity in the sub-RPE and choriocapillaris regions than YN eyes. **(D)** Counting of acellular capillaries revealed that OD eyes have a significantly (p<0.001) higher degree of choriocapillaris atrophy in the macula than YN and ON eyes.

### Characterization of Isolated Monkey Choroidal ECs

Choroidal ECs isolated from the macular region of YN, ON, and OD monkey eyes and cultured in endothelial-selective growth medium exhibit typical colony morphology after seeding (Fig. 2A). Further, our RT-qPCR and flow cytometry measurements revealed that these ECs also express the key markers of endothelial lineage viz. VE-Cadherin (Fig. 2B), VEGFR2 (Fig. 2C), CD146 (Fig. 2D), and CD31 (Fig. 2E). Consistent with their endothelial phenotype, the choroidal ECs also assembled into capillary-like networks when plated on basement membrane hydrogel (*Suppl. Fig*.*1*), a distinctive feature of ECs. Finally, we confirmed that the EC cultures are devoid of RPE cells, as judged by undetectable levels of RPE65 mRNA, a phenotypic hallmark of RPE (*Suppl. Fig*.*2*).

**Figure 2:**
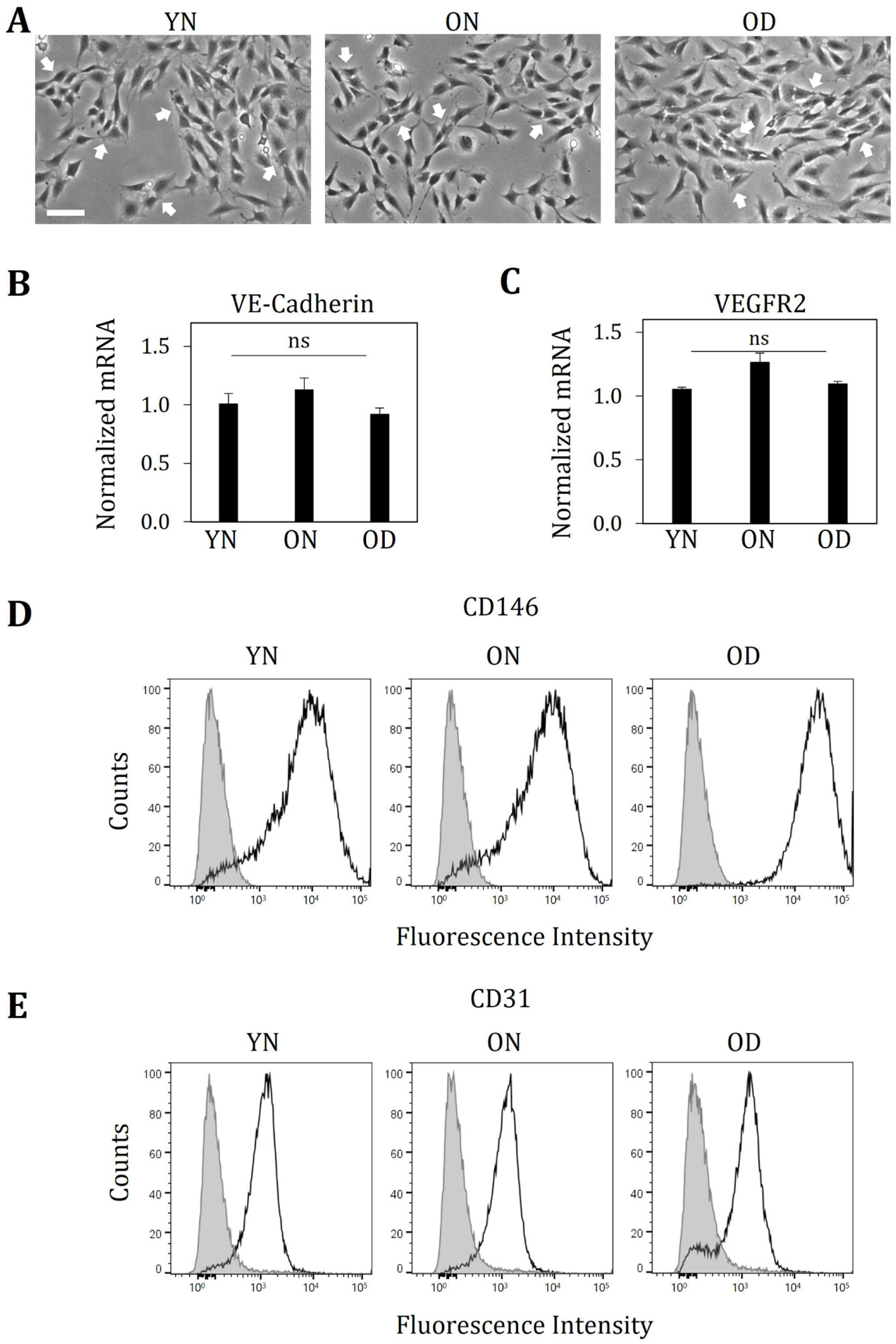
Phenotypic characterization of isolated monkey choroidal ECs. **(A)** Phase contrast images of macular choroidal ECs isolated from YN, ON, and OD monkeys reveal colony formation (arrows) characteristic of EC cultures. Scale bar: 100 um. **(B, C)** Quantitative PCR analysis of the isolated choroidal ECs indicate comparable mRNA levels of VE-cadherin and VEGFR2, key endothelial-specific markers. mRNA levels were normalized w.r.t. GAPDH and expressed as average ± standard error of mean (SEM). **(D, E)** Monkey choroidal ECs were labeled with anti-CD146 or anti-CD31 (empty histograms) antibody, or isotype-matched control antibody (solid gray histogram) and subjected to flow cytometry analysis. Histograms of cell counts versus fluorescence intensity indicate notable expression of the endothelial cell surface markers CD146 and CD31 in these cells.

### Choroidal ECs from Drusen-laden OD Eyes Exhibit Increased Susceptibility to Complement Injury

To assess their response to complement activation associated with AMD, choroidal ECs from YN, ON, and OD monkey eyes were treated with normal human serum (NHS), which activates the alternative complement pathway and promotes membrane MAC deposition [10,18]. Phalloidin labeling of NHS-treated cells revealed that OD cells exhibit the highest degree of cellular injury in response to complement activation, as judged by the extensive disruption of F-actin microfilaments in these cells (Fig. 3A). Quantification of cellular injury showed that OD cells exhibit 1.4-fold (p<0.01) and 2.7-fold (p<0.001) increase in complement susceptibility when compared with ON and YN cells, respectively. Interestingly, this preferential injury of OD cells did not appear to result from higher levels of membrane MAC (C5b-9) deposition, which was found to be comparable across the different conditions (Fig. 3B).

**Figure 3:**
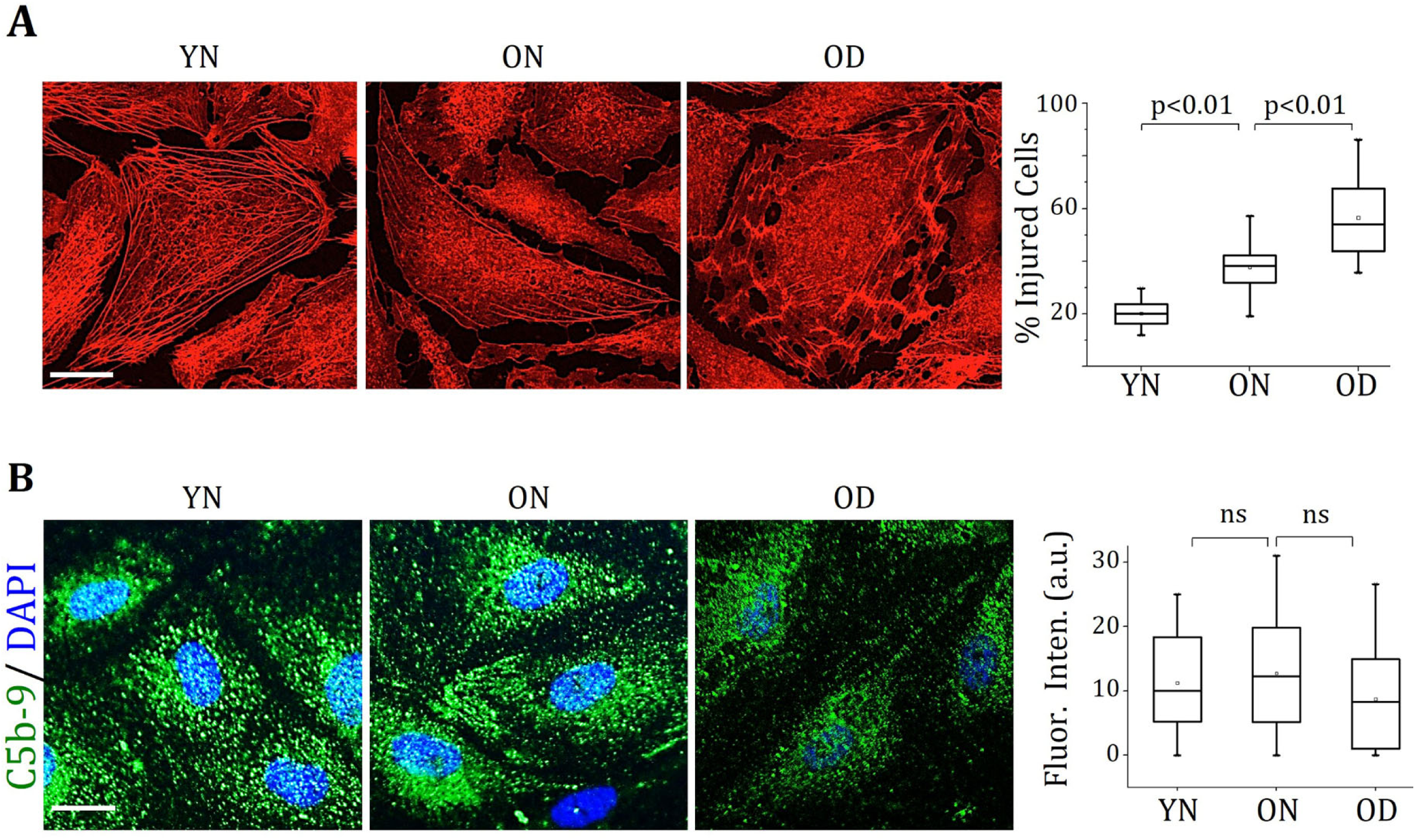
Choroidal ECs from drusen-laden eyes exhibit increased susceptibility to complement injury. **(A)** Choroidal ECs were treated with complement-competent normal human serum (NHS, 5% v/v; 3h, 37°C) prior to phalloidin labeling of F-actin cytoskeletal filaments. Representative fluorescent images and quantitative analysis of F-actin integrity revealed that OD ECs undergo significantly greater cytoskeletal damage when compared with YN and ON cells (P < 0.01; n ≥ 300 cells/condition). Scale bar: 25 um. **(B)** NHS-treated choroidal ECs were labeled with anti-C5b-9 to detect surface membrane attack complex (MAC) deposition. Representative confocal fluorescence images and quantitative image analysis (bar graph; n ≥ 75 cells/condition) reveal no significant differences (P > 0.05) in MAC deposition between these cells after 3h NHS treatment.

### Choroidal ECs from OD Eyes Exhibit Higher Stiffness and Peripheral Actin Organization

Our previous study revealed that senescent RF/6A chorioretinal cells exhibit higher stiffness that, in turn, increases their susceptibility to complement injury [10]. Since cellular senescence is a hallmark of aging, we here asked whether the greater complement injury of OD ECs was associated with age-related stiffening of these cells [14]. Our AFM measurements revealed that, indeed, the OD ECs from drusen-laden eyes exhibit the highest stiffness among the three groups of choroidal ECs (Fig. 4A). Specifically, the median stiffness of OD ECs was found to be two-fold higher (p<0.01) than that of their younger counterparts (YN ECs) and 1.4-fold higher (albeit not significant) than their age-matched controls (ON ECs). Further, consistent with the more pronounced actin cytoskeletal injury seen in NHS-treated OD ECs (Fig. 3A), these cells exhibited the greatest loss in median stiffness (by 78%; p<0.001) in response to complement injury (*Suppl. Fig*.*3*); in contrast, the more resilient ON and YN ECs exhibited only 37% (p<0.01) and 20% (p>0.05) reduction in their stiffness, respectively, upon NHS treatment.

**Figure 4:**
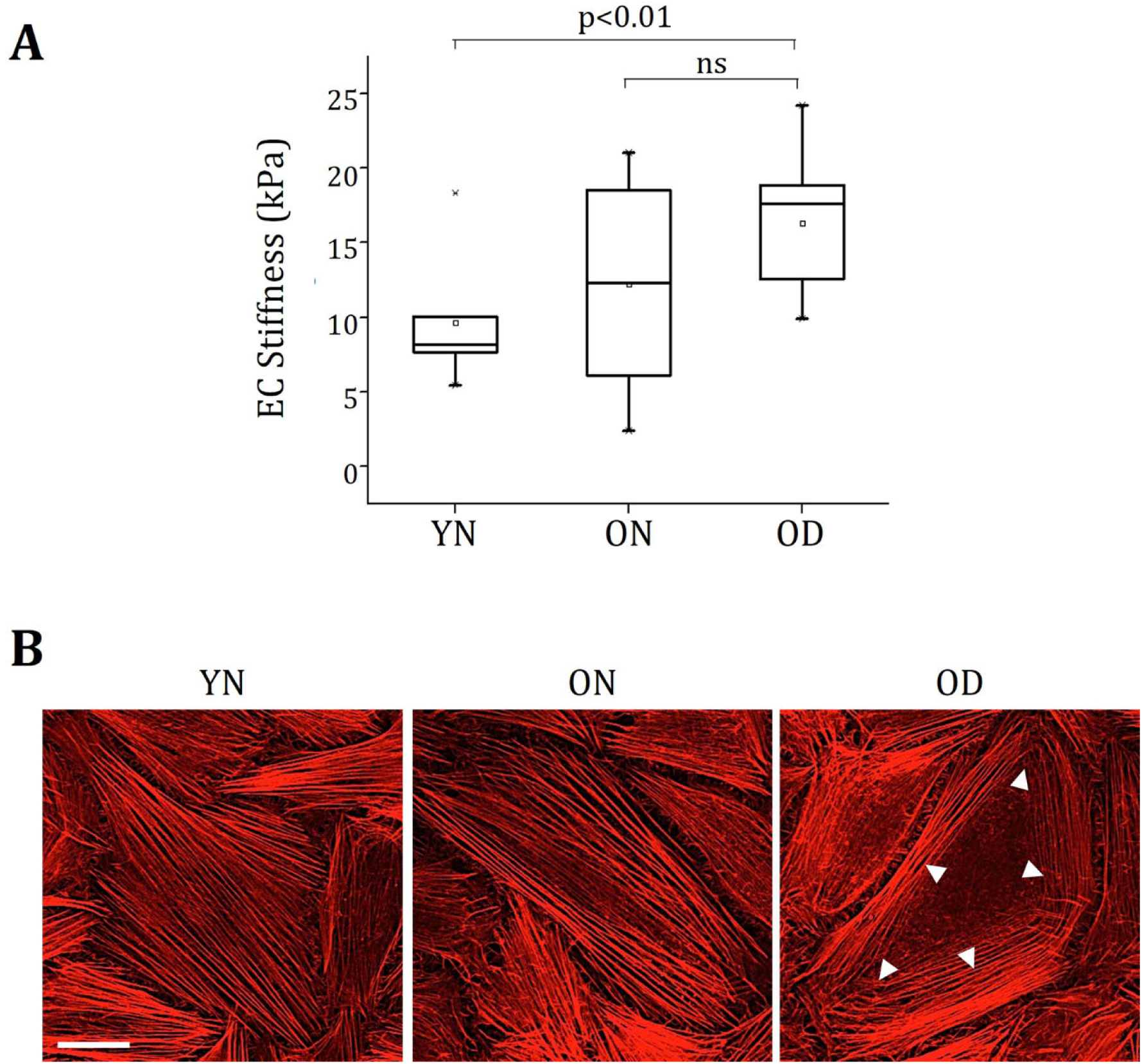
Choroidal ECs from drusen-laden eyes exhibit higher stiffness. **(A)** EC stiffness was measured using a biological-grade AFM fitted with a silicon nitride cantilever tip that was modified with a 5 μm-diameter glass bead. Quantitative analysis of multiple (n≥60) force indentation measurements revealed that the stiffness of choroidal ECs isolated from OD eyes is 2-fold higher than that of their young normal (YN) counterparts (P < 0.05). **(B)** Choroidal ECs were stained with fluorescently-labeled phalloidin to visualize F-actin cytoskeletal filaments. Representative fluorescent images revealed robust longitudinal actin stress fibers in YN and ON ECs but a peripheral actin arrangement in OD ECs, as indicated by the white arrowheads. Scale bar: 50 um.

We and others have previously shown that stiffer cells exhibit a higher density of tension-bearing actin microfilaments [10,19,20]. Interestingly, phalloidin labeling of the choroidal ECs revealed that the stiffer OD ECs exhibit a peripheral actin organization that is markedly distinct from the longitudinal stress fibers observed in YN and ON ECs (Fig. 4B).

### Rac-dependent Cell Stiffening Contributes to Complement Injury of OD ECs

Rac1, a member of the Rho family of small GTPases, promotes peripheral actin organization that, in turn, contributes to cell stiffness [21,22]. Indeed, our G-LISA measurements revealed that the stiffer OD ECs with peripheral actin exhibit markedly higher (∼1.35-fold) levels of Rac1 activity when compared with levels in YN and ON cells (Fig. 5A). Consistent with these findings, treatment of OD cells with a pharmacological Rac1 inhibitor (NSC23776) caused disruption of peripheral actin (Fig. 5B) and a concomitant 2.4-fold decrease (p<0.001) in cell stiffness (Fig. 5C). That Rac1-dependent cell stiffening contributes to the complement injury of OD ECs was confirmed when Rac1 inhibition (with NSC23776) alone resulted in a 2.5-fold decrease (p<0.001) in the number of complement-injured OD ECs (Fig. 5D).

**Figure 5:**
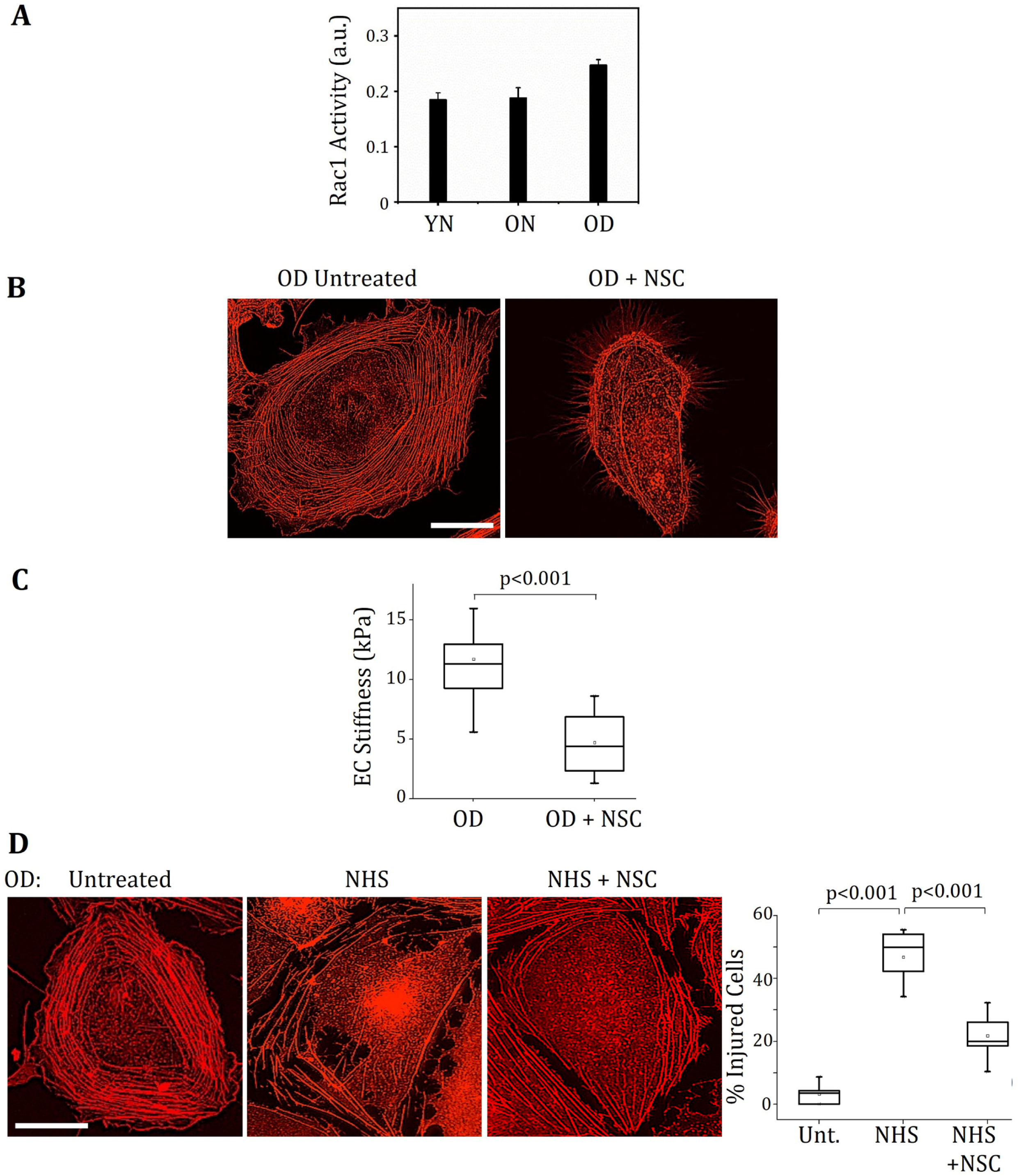
Inhibiting Rac-mediated cell stiffness reduces the susceptibility of OD ECs to complement injury. **(A)** Rac activity in choroidal ECs was measured using Rac1 G-LISA activation assay, which revealed a ∼35% increase in Rac activity in OD ECs when compared with YN cells. Bars indicate average **±** SEM. **(B)** Phalloidin staining of OD ECs treated with pharmacological Rac1 inhibitor (NSC23776; 1 mM) shows disruption of peripheral actin by Rac1 inhibition. **(C)** AFM was used to measure the stiffness of OD ECs treated with or without NSC23776. Quantitative analysis of multiple (n≥60) force indentation measurements indicates that Rac1 inhibition significantly reduces cell stiffness. **(D)** Representative fluorescent images of phalloidin-labeled OD ECs and quantitative analysis of F-actin integrity (box plot; n>100 cells/condition) show that inhibiting Rac-mediated cell stiffness significantly prevents complement injury in OD ECs. Scale bar: 25 um.

### Higher Rho/ROCK Activity in Young ECs Confers Protection from Complement Injury

Since YN ECs with longitudinal actin stress fibers also undergo reduced complement injury, we asked whether the actin organization somehow confers protection from complement injury. To address this question, we looked at the role of small GTPase RhoA that, via its downstream effector Rho-associated kinase (ROCK), generates actin stress fibers via an increase in myosin-dependent cytoskeletal tension (contractility) [12,23,24]. Our Rho-GLISA measurements revealed that, indeed, YN ECs exhibit the highest level of Rho activity, which was ∼1.7-fold higher (p<0.05) than that in OD cells (Fig. 6A). This trend in Rho activation correlated with levels of myosin activity (phosphorylation) in these cells, with YN ECs exhibiting a ∼2-fold higher level (p<0.001) than that in OD cells (Fig. 6B). Importantly, Rho/ROCK-dependent stress fiber organization in YN ECs contributes to their reduced complement injury because treatment of these cells with the pharmacological ROCK inhibitor Y27632 markedly disrupted their actin stress fibers (Fig. 6C) and resulted in a concomitant 4-fold increase (p<0.001) in complement injury of YN ECs (Fig. 6D).

**Figure 6:**
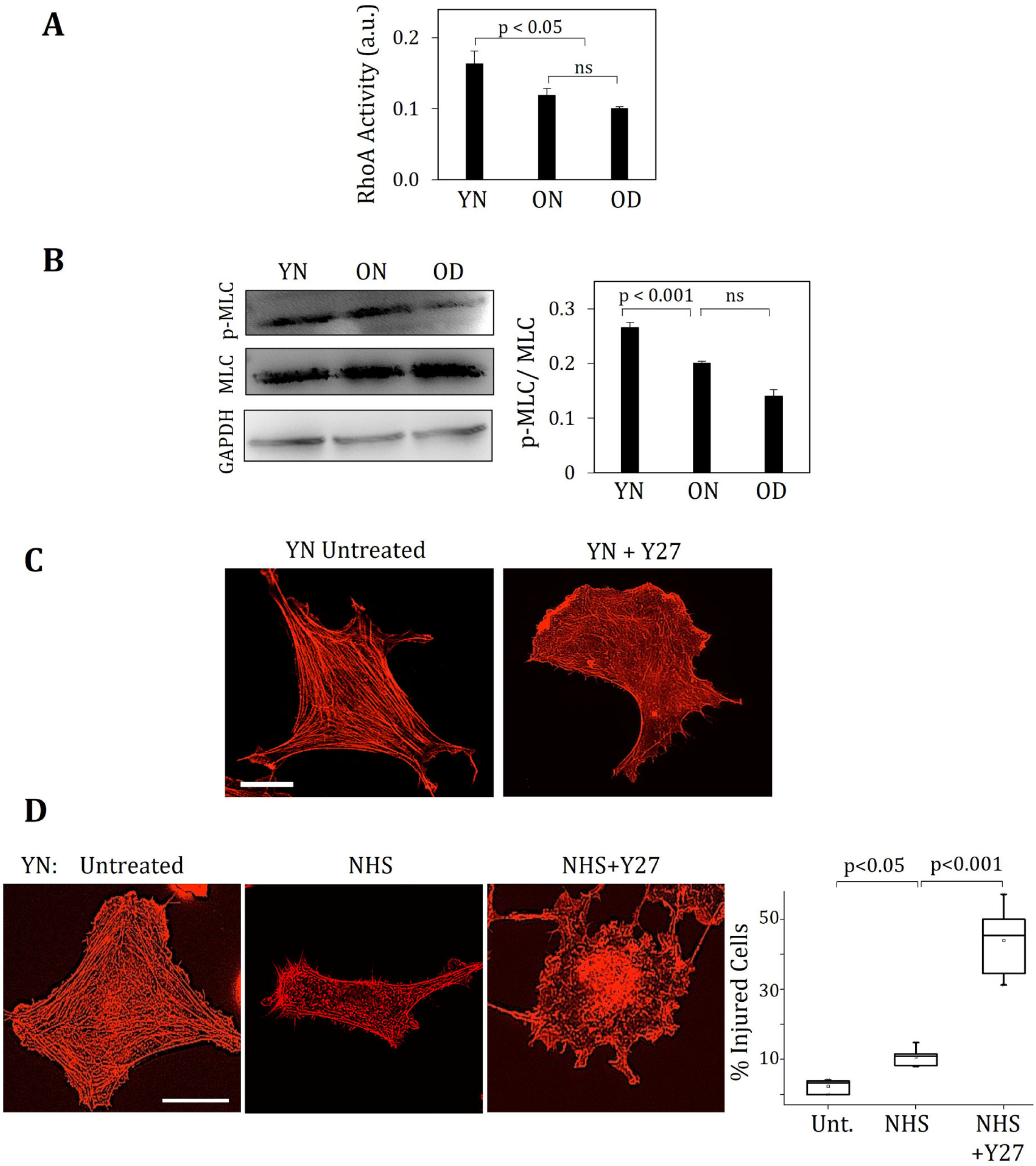
RhoA confers YN ECs protection against complement injury. **(A)** RhoA activity in choroidal ECs was measured using RhoA G-LISA activation assay, which revealed a ∼60% decrease in Rho activity in OD ECs when compared with YN cells (P < 0.05). Bars indicate average **±** SEM. **(B)** Representative Western blot bands (18 kDa) for phosphorylated MLC (pMLC) and their densitometric analysis (mean **±** std. dev.) normalized to total MLC shows that MLC phosphorylation is reduced by ∼50% in OD ECs (P < 0.001). **(C)** Phalloidin staining of YN ECs treated with pharmacological ROCK inhibitor (Y27632; 100 uM) shows disruption of longitudinal actin stress fibers by RhoA/ROCK inhibition **(D)** YN ECs were co-treated with NHS and Rho/ROCK inhibitor Y27632 (100 uM). Representative fluorescent images of phalloidin-labeled cells and quantitative analysis of F-actin integrity (box plot; n>100 cells/condition) show that inhibiting RhoA/ROCK-mediated contractility causes an ∼40% increase in complement injury in YN ECs (P < 0.001). Scale bar: 25 um.

## Discussion

Despite the growing recognition that choroidal vascular degeneration might contribute to AMD progression, the mechanisms underlying choroidal EC loss in early AMD remain poorly understood. In the current study, we used a rhesus monkey model of early AMD that, our findings reveal, exhibits MAC deposition and choriocapillaris atrophy similar to that seen in human AMD eyes. We further show that choroidal ECs isolated from drusen-laden monkeys exhibit increased cytoskeleton-dependent stiffness that, in turn, increases their susceptibility to complement injury. Thus, by demonstrating the translational relevance of rhesus monkeys for the study of choroidal vascular degeneration and identifying EC stiffening as an important determinant, this study strengthens the rationale to more closely examine rhesus monkeys and mechanobiological mechanisms for a deeper understanding of AMD pathogenesis.

Besides forming blood vessels to metabolically support the outer retina, choroidal ECs also appear to provide trophic factors for RPE barrier function and basement membrane remodeling during development [6]. The choriocapillaris is also implicated in the clearance of basal RPE deposits [4]. Thus, the degeneration of such a multifunctional vascular unit may contribute to RPE and photoreceptor defects. This idea has led to a growing interest in understanding how choroidal vessels degenerate in early AMD. One potential mechanism might involve MAC. This terminal product of complement activation is abundantly found around degenerating choroidal vessels (particularly the choriocapillaris) of human early AMD eyes [8,9]. Since MAC can form membrane pores and cause cell lysis, it may promote choroidal EC death and subsequent choroidal vascular loss. However, no mechanistic study has yet examined this possibility in early AMD. This is primarily due to a lack of known animal models that closely recapitulate both the human retinal structure (presence of macula) and aforementioned choroidal vascular phenotype of early AMD [15,25]. As rhesus monkeys resemble humans with regard to retinal structure, AMD susceptibility genes ARMS2 and HTRA1, and age-related drusen phenotype, here we examined their macula for evidence of choroidal vascular changes.

Our rhesus monkey retinal sections revealed significantly higher levels of choriocapillaris-associated MAC immunoreactivity and EC loss in drusen-laden early AMD eyes than in young or old normal (age-matched control) eyes, a trend that is also observed in humans [4,5,8]. Further, our observation that young monkey eyes also exhibit notable MAC deposition, specifically in Bruch’s membrane, is consistent with findings in young human eyes [8]. Thus, these data support the view that while excessive complement activation associated with aging or AMD may contribute to choroidal vascular loss, its low-to-moderate levels in young eyes may have a beneficial role, perhaps in opsonizing sub-RPE debris for clearance by phagocytic cells. Notably, these findings also establish rhesus monkeys as a unique and clinically relevant model for investigating choroidal vascular changes in early AMD.

The observation that MAC deposition increases around the degenerating choroidal vessels of early AMD eyes points towards a potential role of complement activation in choroidal vascular loss. Indeed, treatment of macular choroidal ECs isolated from YN, ON, and OD monkey eyes with complement-competent serum resulted in actin cytoskeletal injury that increased significantly in ON and OD cells. To our knowledge, these findings, which align with previous reports of complement-mediated actin disruption in glomerular epithelial cells [26], are the first to identify a functional difference between choroidal ECs from young, aging, and AMD eyes. These data are also consistent with our previous findings that senescence, a key age-related factor that is implicated in many aging disorders, increases complement injury of RF/6A chorioretinal cells [10]. Notably, the preferential injury of OD ECs was not associated with increased MAC deposition because similar levels of MAC immunoreactivity were found on EC membranes from all three groups of monkeys. Although this observation contradicts the higher MAC levels found in the choroidal sections of ON and OD monkeys, we reason that this difference in MAC levels between EC cultures and choroidal vessels in retinal sections likely results from the prolonged duration of MAC deposition that occurs in aged monkey choroids.

Our AFM measurements show that the more susceptible OD ECs are also significantly stiffer than their younger counterparts. Although this is the first report of choroidal EC stiffening in early AMD, others have reported age-related increase in the stiffness of retinal vessels and non-ocular arteries [27,28]. Furthermore, vascular stiffening often occurs in regions of chronic inflammation such as atherosclerotic lesions where the stiffer ECs exhibit increased sensitivity to inflammatory factors through the process of mechanotransduction [13,29-32]. Thus, our current findings consolidate the growing view that implicates mechanical cues as a key determinant of vascular inflammation [33-35]. Notably, our findings also revealed that while there was no significant difference between the stiffness of OD (AMD) and ON (non-AMD) ECs, the OD cells underwent significantly greater complement injury. Thus, differences in non-mechanical (biochemical) factors, such as expression of membrane-based complement regulators (CD59, CD55, CD46) [4], also likely contribute to the greater complement susceptibility of OD ECs.

Interestingly, these choroidal ECs also differ markedly with regards to their actin cytoskeletal organization, with the OD cells exhibiting robust peripheral actin that is distinct from the longitudinal stress fibers seen in YN ECs. Not surprisingly, we found that this differential actin organization is caused by differences in the activity of their cytoskeletal regulators Rac and Rho/ROCK, members of the Rho family of small GTPases that stabilize peripheral actin and longitudinal stress fibers, respectively [10,12,21,22]. That high Rac contributes to higher stiffness and complement susceptibility of OD ECs was confirmed when disruption of Rac-dependent peripheral actin organization resulted in simultaneous reduction of cell stiffness and complement injury in these cells. Conversely, the relatively higher Rho/ROCK activity in YN ECs appear to protect them from complement activation because ROCK inhibition alone exacerbated complement injury in these cells. Interestingly, this finding contradicts our past observation where Rho/ROCK inhibition in senescent cells reduced complement injury, and vice versa [10]. One possible explanation for this contradiction is that Rho/ROCK regulates complement susceptibility in a biphasic manner where excessively high levels, as might be the case with senescent cells, increases complement injury while the presumably optimal levels in young choroidal ECs exerts protective effects. Consistent with this view, other studies have also reported biphasic inflammatory effects of Rho on ECs wherein Rho/ROCK biphasically regulated endothelial ICAM-1 expression/clustering and EC junctional stability that, together, result in leukocyte-EC adhesion and vascular hyperpermeability [30,36].

In summary, by identifying cell stiffness and its cytoskeletal regulators Rac and Rho as key determinants of choroidal EC susceptibility to complement injury, the current findings offer a new mechanistic insight into choroidal EC loss and CC degeneration associated with early AMD. Importantly, they also confirm the relevance of the rhesus monkey model for future studies that more deeply examine the mechanobiological mechanisms by which choroidal EC stiffness mediates this deleterious effect. Whether such mechanisms alter the cell’s ability to ‘prevent’ or ‘repair’ membrane injury via expression of complement regulatory factors or endocytosis/exocytosis, respectively [37-39], also remains to be seen. Ultimately, use of the rhesus monkey model of early AMD to uncover the mechanobiological basis for complement-mediated choroidal EC loss may help identify new classes of therapeutic targets for effective AMD management in the future. These efforts can dovetail with other stiffness-normalization approaches that are currently being developed by the pharmaceutical industry for various debilitating complications [40].

## Acknowledgements

Acknowledgement is made to the donors of Macular Degeneration Research, a program of BrightFocus Foundation, for support of this research (Grant M2016161 to K.G.). This work was also supported by the Initial Complement Funds provided by the BCOE at UC Riverside (to K.G.), Start-up Funds provided by the Doheny Eye Institute (to K.G.), The Stephen Ryan Initiative for Macular Research (RIMR) Special Grant from W.M. Keck Foundation (to Doheny Eye Institute), an Unrestricted Grant from Research to Prevent Blindness, Inc. (to K.G. and UCLA Ophthalmology), and NIH grants 51OD011092 and S10RR024585 (to ONPRC), and P30EY010572 (to Casey Eye Institute).

## Author contributions statement

A.P.C. designed and performed experiments, analyzed data, and wrote the manuscript; J.S. performed experiments, I.S.T. and N.M. performed experiments and analyzed data, M.A. and N.P. analyzed data, L.R. performed experiments, M.N. and T.M. contributed new reagents, and K.G. conceived the idea, designed experiments, analyzed data, and wrote the manuscript. All authors reviewed and edited the manuscript.

## List of Supplementary Material Online

I. **Materials and Methods –** Cell Isolation
II. **Figures and Legends–**

**Fig. 1:** Isolated choroidal ECs form capillary-like networks in vitro

**Fig. 2:** Choroidal EC culture is devoid of RPE cells

**Fig. 3:** Choroidal ECs from drusen-laden OD eyes undergo marked softening in response to complement injury

