## Supporting Information for "Cell Stiffening Contributes to Complement-mediated Injury of Choroidal Endothelial Cells in Early AMD"

**Supplementary Materials and Methods**

**Cell Isolation**

Macular choroidal endothelial cells (ECs) were isolated from three groups of female rhesus monkeys viz. young normal (YN; 6 years old), old normal (ON; 20 years old), and old monkeys with severe drusen (OD; 19 years old). Specifically, eyes were harvested from monkeys undergoing necropsy, and after removal of exterior tissue, were placed in 10% betadine (Fisher Scientific) for 15 minutes at room temperature. Eyes were washed three times with HBSS prior to dissecting off the anterior chamber. Four radial relief cuts were made to the retinal/scleral tissue before placing it in isolation buffer composed of MCDB131 basal medium (Corning) containing 30 mM HEPES and 1x antibiotic/antimycotic (Life Technologies). Without traversing the sclera, 8 mm punch biopsies were obtained from the central region containing the inner choroid and choriocapillaris/Bruch's membrane complex. The neural retina was removed, leaving the biopsy punch in place. Isolation buffer was replaced with 0.1% BSA in HBSS and retinal pigment epithelial (RPE) cells were gently brushed off. The tissue was washed several times with 0.1% BSA until RPE was completely removed. Next, the biopsy punches were detached from periphery and the choroid was removed from the sclera prior to washing three times with 0.1% BSA in HBSS. Choroidal tissue was spun down and transferred to a culture plate containing isolation buffer prior to cutting into 1 mm pieces. Choroidal tissue was collected and washed two times with HBSS prior to incubating with 0.5% trypsin (15 min, 37$^{o}$C). Next, trypsin was removed and choroidal tissue was incubated with isolation buffer containing DNAse (2.7 Kunitz/ul stock; Qiagen) and 0.1% collagenase (Sigma) on a rocker, vigorously shaking every 5 min until tissue was digested (30 min, 37$^{o}$C). Suspension was then washed three times with 0.1% BSA HBBS prior to being filtered through a sterile 70 μm mesh followed by rinsing with 0.1% BSA in HBSS. Cell suspension was spun down and resuspended in enriched endothelial growth medium composed of MCDB131 medium supplemented with fetal bovine serum (FBS; 15%; HyClone GE), Glutamax (2mM; Life Technologies), vascular endothelial growth factor (VEGF; 1 ng/ml; Biovision), epidermal growth factor (EGF; 5 ng/ml; Sigma), basic fibroblast growth factor (bFGF; 8 ng/ml; Life Technologies), heparin (100 µg/ml; Sigma), ascorbic acid (50 µg/ml; Sigma), 1x antibiotic/antimycotic prior to seeding onto fibronectin-coated (1 µg/cm$^2$; Corning) tissue culture dishes. Four hours after cell plating, cells were rinsed with PBS to remove non-adherent cells and supplemented with fresh endothelial growth medium. Cells were grown on fibronectin-coated dishes for the first two passages and, thereafter, subcultured on gelatin-coated (0.5% w/v) dishes, with medium replaced every two days.

Retinal pigment epithelial (RPE) cells were isolated from an infant rhesus monkey undergoing necropsy. Eyes were cut open and without traversing the sclera, the neural retina was removed. The retinal pigmented epithelial (RPE) cells were gently brushed off from the Bruch's membrane and collected. RPE cells were spun down and frozen in RNAlater stabilization solution (Invitrogen) for subsequent PCR studies.

**Supplementary Figure Legends**

**Supplementary Figure 1: Isolated choroidal ECs form capillary-like networks in vitro**

Phase contrast images of the monkey choroidal ECs plated on basement membrane extract for 6h reveal their ability to spontaneously form capillary-like networks, a hallmark of endothelial cells. Scale bar: 100 um.

**Supplementary Figure 2: Choroidal EC culture is devoid of RPE cells**

Quantitative PCR analysis of 10 day-old macular choroidal EC cultures probed for RPE65 mRNA, a key RPE-specific marker, indicate a lack of RPE cell contamination. RPE cells from an infant rhesus monkey were used as positive control. mRNA levels are normalized w.r.t. GAPDH and expressed as average ± SEM.

**Supplementary Figure 3: Choroidal ECs from drusen-laden (OD) eyes undergo marked softening in response to complement injury**

Stiffness of untreated or NHS-treated ECs was measured using a biological-grade AFM fitted with a pre-calibrated probe tip of 70 nm radius. Quantitative analysis of multiple (n=45) force indentation curves revealed that NHS treatment leads to a significantly greater reduction (by 78%; $p<0.001$) in the median stiffness of OD ECs than that of ON and YN ECs (37% and 20% reduction, respectively).

### Suppl. Fig. 1

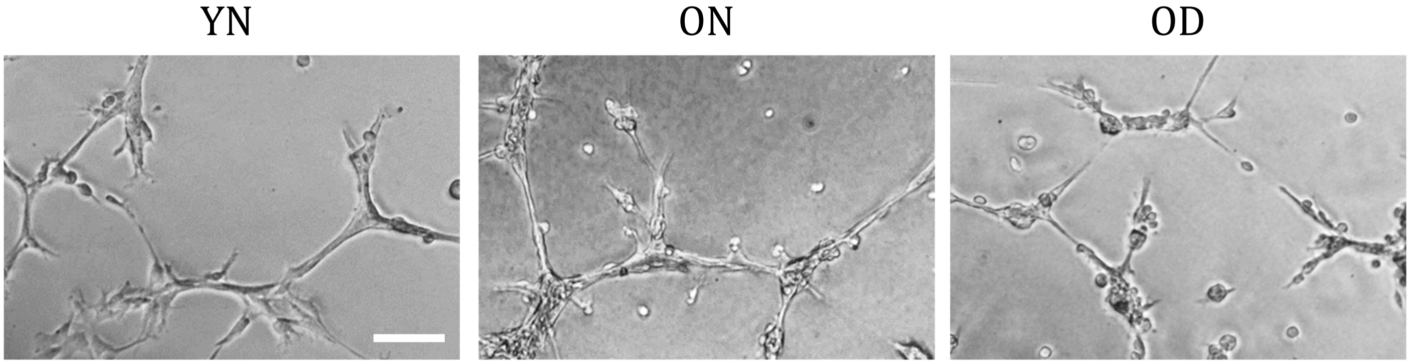

**Suppl. Fig. 2**

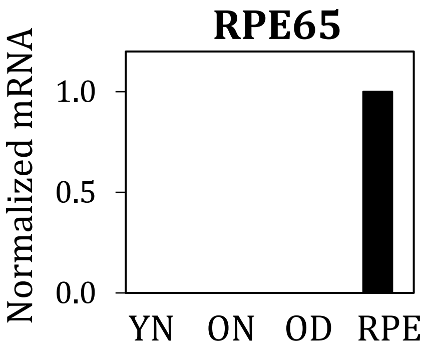

**Suppl. Fig. 3**

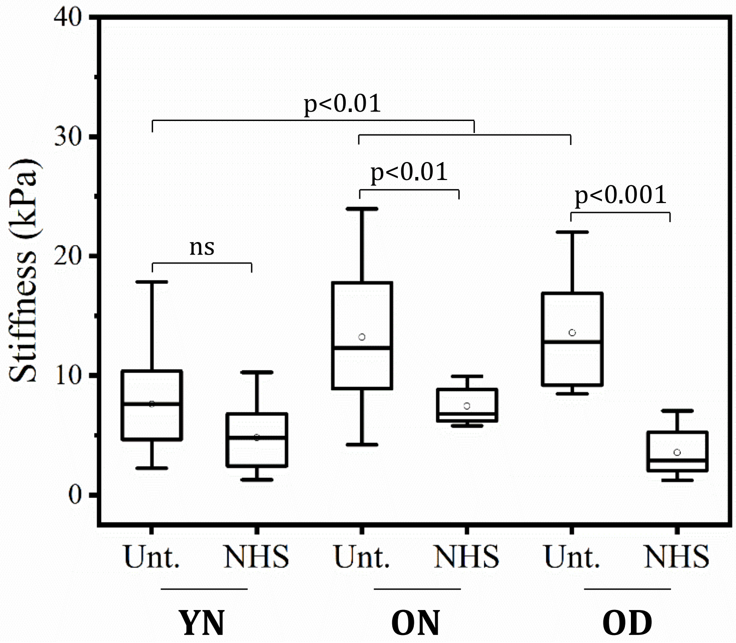

40
30
20
10
0
Stiffness (kPa)
p<0.01
p<0.01
p<0.001
ns
Unt.
NHS
Unt.
NHS
Unt.
NHS
YN
ON
OD
